# Inferring systemic metabolic and oxidative stress susceptibility in normal-tension glaucoma through targeted skin fibroblast analysis

**DOI:** 10.1101/2025.09.21.677625

**Authors:** Alexander von Spreckelsen, Katarina Stoklund Dittlau, Sarkis Saruhanian, Rupali Vohra, Arevak Saruhanian, Blanca I. Aldana, Kristine Freude, Miriam Kolko

## Abstract

**Aim:** Enhance the understanding of intrinsic metabolic and oxidative stress vulnerability in normal-tension glaucoma (NTG) pathophysiology by targeted profiling of skin fibroblasts from NTG and control donors.

**Background:** NTG is a primary open-angle glaucoma subtype characterised by glaucomatous neurodegeneration without elevated intraocular pressure. Increasing evidence links systemic metabolic and oxidative stress vulnerability to NTG pathophysiology, making non-ocular, somatic cells promising surrogate systems for assessing neurodegenerative predisposing mechanisms.

**Methods:** Skin fibroblast cultures were obtained from four female NTG and four age- and gender-matched control donors. Targeted metabolic profiling of mitochondrial function, glycolytic capacity, and glucose and amino acid metabolism was performed using the Seahorse assay, gas chromatography-mass spectrometry, and high-performance liquid chromatography. Oxidative stress resilience to hydrogen peroxide was assessed employing the lactate dehydrogenase release assay.

**Results:** Pure skin fibroblast cultures were obtained for all included donors. NTG and control fibroblast exhibited similar mitochondrial and glycolytic function. No group difference was demonstrated in relative glucose metabolism and absolute amino acid profile. Control and NTG fibroblasts exhibited similar susceptibility to oxidative stress.

**Conclusion:** NTG skin fibroblasts exhibit similar mitochondrial and glycolytic function, glucose metabolisation, amino acid profile, and oxidative stress resiliency compared to controls. Future studies should focus on mapping cell- and tissue-specific differences through combined genetic and transcriptomic profiling to guide and stratify the functional assessment of different endotypes within the NTG disease spectrum. Cell-specific dysregulation and the need for individual mechanistic grouping diminish the applicability of skin fibroblasts as a model system for exploiting NTG pathophysiology.

## Introduction

Glaucoma is a heterogeneous group of optic neuropathies characterised by progressive dysfunction and loss of retinal ganglion cells (RGCs) [1]. Primary open-angle glaucoma (POAG) is the most common glaucoma subtype [2], which is further divided into normal-tension glaucoma (NTG) and high-tension glaucoma, separated on the basis of the absence or presence of clinically defined elevated intraocular pressure (IOP) [1]. More than 80 million people are affected by POAG worldwide [2], while the current extent of undiagnosed cases is estimated to surpass this number [2]. Definitive causes remain elusive; however, age, familial disposition, and IOP are well-established clinical risk factors, with the latter being the only currently therapeutically modifiable target [1]. Irrespective of the POAG phenotype, lowering IOP halts glaucoma incidence and progression [3, 4]. However, more than 40% of high-tension glaucoma patients [3] and 10% of NTG patients still progress despite a baseline IOP reduction of 25% or more [4]. As such, due to diagnostic delay and therapeutic insufficiency, the lifetime risk of blindness is significant [5] and has in some studies been demonstrated as high as 42% and 16% for developing uni- and binocular blindness, respectively [6]. To counteract the increasing glaucoma burden, substantial advancements within prophylaxis, diagnostics and personalised medicine are needed [2].

NTG constitutes up to half of all POAG cases [7], with a relative overrepresentation of females and people of Asian descent [7]. Like many neurodegenerative diseases, NTG is considered a multifactorial disease, with major pathophysiological hypotheses focusing on diminished IOP tolerance [8], excessive IOP fluctuations [8], vascular dysregulation [9, 10], abnormal glymphatics [8, 11], reactive neuroinflammation [12], as well as bioenergetic crisis [13] and oxidative stress [14]. Especially bioenergetic metabolism and mitochondrial function play a central role in the RGCs’ ability to cope with stress [13, 15–17]. Due to the complex and multi-compartmentalised anatomy and physiology of the RGCs, these cells require tremendous energy and support demands to sustain their optimal function [13]. These requirements also render RGCs particularly predisposed to disease, consequently highlighting increasing evidence of local and systemic metabolic dysfunction as key players in glaucoma development and progression [13, 18–20]. Locally, RGC mitochondrial abnormalities can already be detected in early-stage glaucomatous retinas [21], i.e. before noticeable RGC loss [21]. In addition, abnormal aqueous humour metabolite profiles [13, 22] and metabolic dysfunction of several other ocular cell types [23–28] support the presence of impaired local metabolic function early in glaucoma pathogenesis. In favour of a broad bioenergetic involvement, systemic metabolic and mitochondrial dysfunction are a common finding in POAG patients [29–32], especially in NTG subjects [32]. Disturbances of several key pathways within essential bioenergetic, redox, amino acid, and lipid metabolism in various fluid compartments (plasma, serum) and peripheral cell types are a common finding in POAG patients [20–22, 32–35], even decades before a POAG diagnosis [35]. Furthermore, worsened disease prognosis is directly associated with the degree of compromised systemic mitochondrial function [32], while low plasma levels of glycolytic and oxidative phosphorylation substrates such as pyruvate, lactate, acetate, and β-hydroxybutyric acid increase the odds of POAG development [35]. Additionally, selective RGC impairment and loss are frequently observed in several mitochondriopathy-related optic neuropathies without extraocular disease [36–38]. Conversely, clinical intervention studies find that mitochondrial substrates maintain RGC function in POAG patients [39, 40], underpinning the importance of metabolic support in maintaining RGC health. As the genetic background underlying systemic mitochondrial and metabolic dysfunction is somatically shared [31, 41–49], these systemic metabolic vulnerabilities may enact a contributing effect rather than being a sole downstream disease consequence. Overall, it is reasonable to assume that *intrinsic* bioenergetic perturbation is a major contributor to glaucoma susceptibility.

Oxidative stress is another hallmark of POAG pathophysiology [50] and refers to a disturbed redox state characterised by excessive pro-oxidation, superseding the capacity of intrinsic cellular antioxidant defence mechanisms [50]. The resulting oxidative damage to DNA, RNA, protein, and lipids augments intrinsic cellular function (e.g., endoplasmic reticulum stress, mitochondrial dysfunction, inflammation, disturbed autophagy), drives mutagenesis, disturbs cellular signalling (e.g., disturbed nitric oxide (NO) mediated vasoregulation), and prompts cell death [51]. Local oxidative stress may facilitate glaucomatous neurodegeneration by directly targeting RGCs or indirectly by modulation of processes in structures vital for maintaining RGC health, including the trabecular meshwork, the lamina cribosa, the optic nerve head, and the retinal microvascular systems [51]. As for metabolic vulnerability in POAG, a systemic oxidative stress component is potentially linked to the disease pathogenesis. POAG patients exhibit diminished antioxidant capacity in serum [52–55] correlating with worsening visual field prognosis [54] and RGC loss [55], while also demonstrating increased levels of oxidative stress damage markers [56, 57]. Analogous deficits have likewise been demonstrated in lymphoblasts of POAG patients, showing enhanced superoxide production resulting from electron loss from complex I of the electron transport chain (ETC) [29]. This combined evidence links glaucoma vulnerability to oxidative stress susceptibility.

NTG exhibits partial IOP-independent glaucomatous neurodegeneration. Several aetiologies are believed to influence NTG development, and these causes may vary across individuals and disease stage [47, 48]. Understanding the pathophysiological mechanisms that drive and counteract IOP-independent retinal neurodegeneration is crucial for improving patient treatment. Systemic glaucoma predisposing mechanisms may be accessible in various systemic cell types [29–32], which makes the potential dysfunction of other somatic cells a promising disease model for addressing mechanisms underlying RGC resilience. Additionally, this approach circumvents the ethical barrier and neurodegenerative bias of examining excised and post-mortem retinal tissue [58]. As NTG is overrepresented in females relative to males [7], female gender is associated with an increased risk of NTG progression [59], and NTG pathophysiology may systematically differ across genders [60–63], this study specifically aims to assess and expand the current knowledge on bioenergetic perturbations and oxidative stress susceptibility in glaucoma pathophysiology through targeted investigation of skin-derived fibroblasts from female NTG patients and control subjects.

## Methods

### Patient recruitment and sample size considerations

This study was approved by the Danish National Committee on Health Research Ethics (project identification code: H-19038704; primary approval: March 5, 2020; extension approval: November 23, 2022) and was conducted in accordance with the principles outlined in the Declaration of Helsinki of 1975. Written informed consent was obtained from each participant for inclusion in the study prior to enrollment. Additionally, participants provided oral consent after the extent of the study was explained to them verbally.

Patient recruitment has previously been described [64]. NTG and control donors were recruited between March 28, 2019, and January 25, 2022. Briefly, age-matched female NTG and control patients were recruited from the ophthalmological departments of Rigshospitalet-Glostrup (Denmark) and Holbaek (Denmark) for glaucoma ambulatory control or presurgical evaluation for cataract, respectively. All patients with NTG underwent a comprehensive eye examination performed by a glaucoma specialist [64]. Controls had a similar eye exam performed by either a glaucoma specialist or a general ophthalmologist. All patients with NTG were examined within three months before inclusion. At least three ophthalmologists agreed on the diagnosis, and six perimetries were performed before inclusion. Patient reports confirmed that IOP measurements had never exceeded 21 mmHg. All patients had reported IOP prior to treatment, and at least six IOP measurements were performed before enrolment in the study.

Inclusion criteria for patients with NTG were as follows: untreated IOP below 21 mmHg at different times of the day (8 a.m. – 5 p.m.); open iridocorneal angles determined by gonioscopy; optic disc cupping characterised by a violated ISNT rule; glaucomatous visual field loss identified by Humphrey or Octopus perimetry, and significant nerve fibre loss detected by ocular coherence tomography. Control patients were required to fulfil the same criteria, except for exhibiting signs of glaucomatous damage.

Exclusion criteria for all participants were as follows: male gender; a medical history of ocular trauma; competing eye conditions other than glaucoma affecting the optic nerve (e.g. optic dis drusen); significant systemic diseases, i.e., dysregulated hypertension, heart failure, hypercholesterolemia, diabetes mellitus, autoimmune diseases, and previous cerebral infarct or haemorrhage; subjects who could not comply with procedures; subjects under 50 years of age, and subjects who smoked at the time of inclusion. Demographic information, medical anamnesis, and general ophthalmological data were collected to check whether participants met the inclusion and exclusion criteria.

Recruited sample sizes were based on preliminary proof-of-concept experiments on differences in mitochondrial respiration between NTG and control skin fibroblasts. To demonstrate a difference in basal mitochondrial respiration of 3000 µmol O2/min/mg protein with a standard deviation of 1500 pmol O_2_/min/mg protein using a two-sided significance level of α = 0.05 and a power of 0.2, a total of four patients per group were included.

### Skin biopsy procedure and fibroblast culture

0.3 mm upper-arm skin biopsies were obtained under sterile conditions using a dermatome (KAI 2517202; Mediq) and local anaesthesia (SAD lidocaine-adrenaline 20mg/5µg/mL). The excised skin biopsy was collected with a sterile tweezer and transferred to a 15 mL Falcon tube containing fibroblast culture medium (1% penicillin/streptomycin (#P0781, Sigma Aldrich), 1% non-essential amino acid solution (#M7145, Sigma Aldrich), 1% GlutaMAX^TM^ Supplement (#35050061, Fisher Scientific), 10% fetal bovine serum (#A5670701, Fisher Scientific), Dulbecco’s Modified Eagles Medium/Nutrient Mixture F-12 Ham (DMEM/F12; #11-330-057, Fisher Scientific)). After transferring into a biosafety cabinet, the skin biopsy was washed with Dulbecco’s phosphate-buffered saline (dPBS; #14190136, Fisher Scientific) and cut into four equal-sized pieces. Each piece was placed into a Leighton tube containing fibroblast culture medium and incubated for two weeks under humidified conditions at 37 °C and 5% CO_2_. Migrated fibroblasts were detached and transferred into T75 flasks for further expansion. For a passage, fibroblasts were washed with dPBS, incubated for 4 min in 0.25% trypsin-EDTA (#25200056; Fisher Scientific), collected with culture medium, and centrifuged at 1200 x RPM for 3 min. The cell pellet was resuspended in culture medium and the cell suspension dispensed into new T-75 flasks. The media was changed the following day and thereafter on a thrice-weekly basis. Fibroblasts were passaged when reaching a confluency of 70-80%. At passage two, fibroblasts were cryopreserved in culture medium containing 20% fetal bovine serum and 10% dimethyl sulfoxide (#D2438, Sigma Aldrich).

Donor fibroblasts at passages three and four were used for the functional experiments in this study. Cell culture and incubation experiments were performed under humidified conditions at 37 °C and 5% CO_2_. Unless stated otherwise, all solutions used for preincubation, rinsing, or treatment of the fibroblasts were dispensed at 37 °C. Phase-contrast images of fibroblasts were obtained using the Invitrogen EVOS XL Core Imaging System (#AMEX1000, Fisher Scientific), equipped with 4x, 10x, 20x, and 40x long-working-distance plan phase infinity-corrected objectives.

### Immunocytochemical fibroblast profiling

Fibroblasts were plated on non-coated 12 mm #1.5 glass coverslips in 4-well plates (#176740, Fisher Scientific) at a density of 10,000 cells/cm^2^ two days prior to fixation. Following a wash with dPBS, fibroblasts were fixed in 4% formaldehyde (#9713.1000, VWR) for 20 min at room temperature. Then, the fibroblasts were washed three times for 5 minutes in dPBS. Subsequently, cells were permeabilised in 0.2% Triton X-100 (#X100, Sigma Aldrich) dPBS (PBST) for 20 min and incubated for 30 min with 5% bovine serum albumin (BSA; #A3059, Sigma Aldrich) in 0.2% PBST. Thereafter, cells were incubated with an anti-vimentin antibody (#18-0052, Invitrogen) diluted 1:500 in 5% BSA in 0.2% PBST at 4 °C overnight. The next day, cells were washed thrice with dPBS for five minutes each and incubated in the dark for one hour with donkey anti-mouse Alexa Fluor™ 488 (#A-21202, Fisher Scientific) diluted 1:1000 in 5% BSA in 0.2% PBST. From this moment, all procedures were performed shielded from light. Then, cells were washed three times in 0.2% PBST for five minutes each, followed by a seven-minute incubation in 0.1% DAPI (#62248, Fisher Scientific) in 0.2% PBST. Then, the cells were washed four times for 5 minutes with 0.2% PBST and mounted on glass object slides containing DAKO fluorescent mounting medium (S302380-2, Agilent Technologies) and allowed to dry for 20 minutes before coverslip sealing with nail polish.

Images were acquired on a Leica DMRB microscope equipped with a DeltaPix camera (#COOL07DPXM) using 16x (PL FLUOTAR 16x/0.50 IMM) and 40x (PL FLUOTAR 40x/1.00-0.50 Oil) objectives and the DeltaPix InSight software. Fibroblast culture purity was assessed by quantifying the Vimentin^+^/DAPI^+^ fraction of positive cells using ImageJ 1.54p software. A minimum of 700 cells per cell line were selected randomly based on DAPI^+^ expression from 10 representative images at 16x magnification.

### Seahorse assay

The mitochondrial function and glycolytic capacity of skin NTG and control fibroblasts were simultaneously assessed using the Seahorse XFe96 Analyser (Agilent Technologies) [65].

Two days before each experiment, fibroblast cell lines were seeded in a Seahorse XFe96/XF Pro cell culture microplate (#103794-100, Agilent Technologies) at a cell density of 180,000 cells/cm^2^. Complete culture media change was performed the following day. The fibroblasts were microscopically monitored daily to ensure cell health and verify uniform monolayer distribution. On the day of the experiment, fibroblasts were washed and incubated in reconstituted and sterile-filtered glucose-, pyruvate-, and glutamine-free DMEM-based assay medium (#d5030, Sigma Aldrich) for one hour under CO_2_-free conditions to facilitate glycogen and glucose depletion. Meanwhile, the seahorse cartridges were loaded with selected respiratory substrates (Port A: Glucose, c_final_ = 6 mM; Port B: Oligomycin A, c_final_ = 1 µM; Port C: Carbonyl cyanide-4 phenylhydrazone (FCCP), c_final_ = 2 µM; Port D: Rotenone and Antimycin A, c_final_ = 1 µM of each) and allowed to calibrate. All respiratory substrates and media were prepared on the day of each experiment, using hydrochloric acid or sodium hydroxide to adjust the pH to 7.4.

A customised template was applied to assess mitochondrial and glycolytic cellular bioenergetics simultaneously. Briefly, each experiment consisted of three baseline readings followed by four consecutive injection periods of three readings (three minutes) spaced by three-minute mixing periods [65].

All mitochondrial parameters were estimated according to the Seahorse Agilent Technologies instructions (Fig 1A). Non-mitochondrial respiration was calculated as the last measured oxygen consumption rate (OCR) value after concurrent rotenone and antimycin A injection. Basal mitochondrial respiration was estimated by subtracting non-mitochondrial respiration from the third post-glucose injection OCR. Maximal respiration was calculated by subtracting non-mitochondrial respiration from the maximal OCR measurement after FCCP injection. Proton leakage was determined as the OCR difference between the minimum OCR after oligomycin A injection and non-mitochondrial respiration. ATP-linked respiration was estimated to be the difference between basal mitochondrial respiration and endogenous proton leakage. Mitochondrial spare respiratory capacity was determined as the difference between the maximal and baseline rates of mitochondrial respiration.

**Fig 1.**
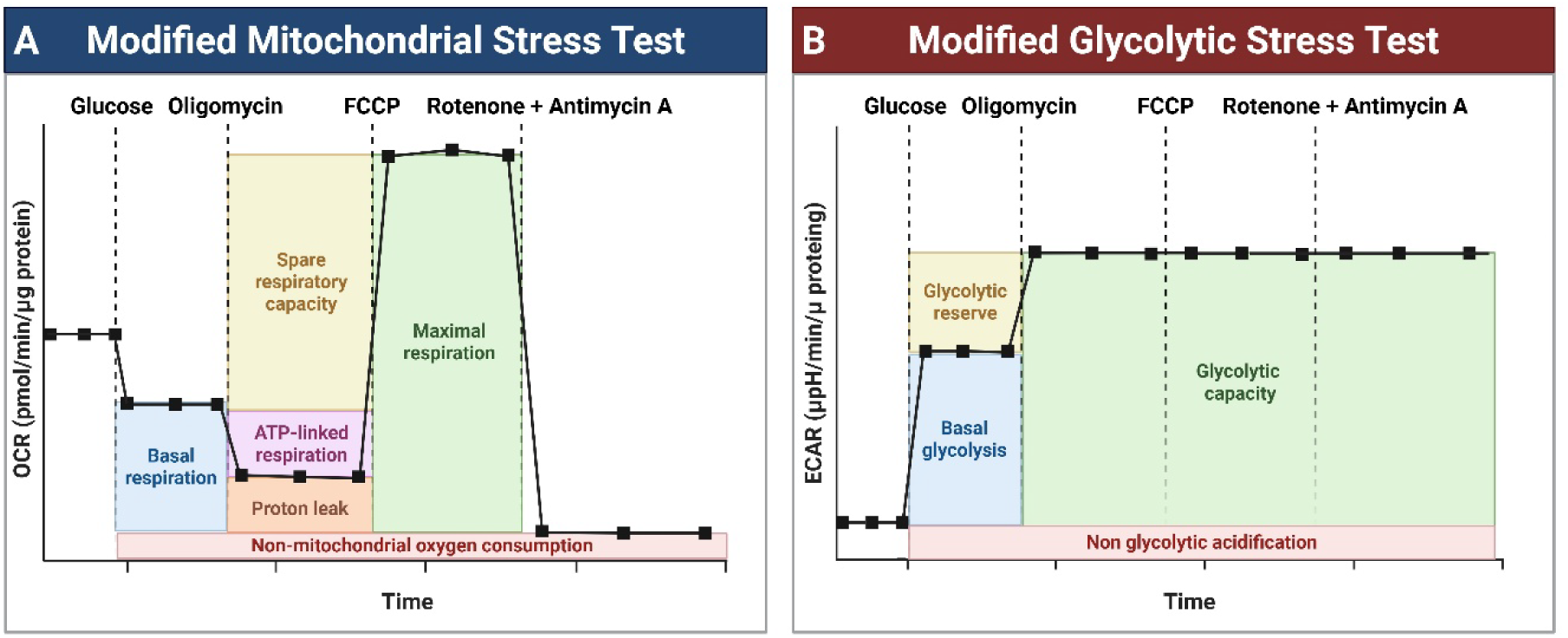
Seahorse template overview: Assessment of (A) mitochondrial and (B) glycolytic function of NTG and control fibroblasts using a modified Seahorse template. All mitochondrial and glycolytic functional parameters can be simultaneously estimated with this setup, except for non-glycolytic acidification. NTG: Normal-Tension Glaucoma; FCCP: Carbonyl Cyanide-4 Phenylhydrazone; OCR: Oxygen Consumption Rate; ECAR: Extracellular Acidification Rate. Created with BioRender.com and adapted from a preformed template by Daisy Shu.

All glycolytic parameters were estimated according to the Seahorse Agilent Technologies instructions (Fig 1B). The basal glycolytic rate was determined as the rate difference between the maximum extracellular acidification rate (ECAR) before oligomycin A injection and the last ECAR measurement before glucose injection. Glycolytic capacity was estimated as the rate difference between the maximum ECAR after oligomycin A injection and the last ECAR measurement before glucose injection. Absolute glycolytic capacity was determined as the ECAR difference between basal and maximal glycolytic capacity.

### Glucose metabolism profiling by stable isotopic labelling and gas chromatography-mass spectrometry

Targeted characterisation of the fibroblast glucose metabolism was performed employing stable isotopic labelling with uniformly ^13^C-labeled glucose ([U-^13^C]glucose) and gas chromatography-mass spectrometry (GCMS) [66, 67].

Two days before each experiment, fibroblasts were seeded in culture medium at a density of 52,000 cells/cm² in 6-well tissue plates (#140675, Fisher Scientific). On the day of the experiment, fibroblasts were washed with 37 °C dPBS and incubated in glucose-, pyruvate-, and glutamine-free culture medium (#A1443001, Sigma Aldrich) supplemented with [U-^13^C]glucose (#CLM-1396-PK; Cambridge Isotope Laboratories) to a final concentration of 6 mM for 4 hours. Afterwards, incubation media were collected and stored at −80 °C until further assessment. Meanwhile, the cells were flushed twice with ice-cold dPBS and submerged in ice-cold RNAse-free ethanol (#11119915001, Sigma Aldrich) for 15 min. Then, cells were detached by manual scraping and collected in Eppendorf tubes at 4 °C (#EP022363344, Sigma Aldrich). The Eppendorf tubes were centrifuged at 20,000 x *g* for 20 minutes at 4 °C, and the supernatants containing the metabolites of interest were collected and stored at −80 °C until lyophilisation. Cell pellets were used for protein normalisation using the BCA assay.

The lyophilised cellular extracts were reconstituted in MilliQ water and centrifuged at 20,000 x *g* for 20 min, at 4 °C. The supernatant was collected for analysis with GCMS and high-pressure liquid chromatography (HPLC). The supernatant was acidified (pH = 1-2) and evaporated under pure nitrogen. Analytes were extracted in an organic phase using 96% ethanol and benzene (c_final_: 1:3 (V/V); #401765, Sigma-Aldrich), and derivatised using dimethylformamide (c_final_: 1:7 (V/V); #270547, Sigma-Aldrich) and N-tert-butyldimethylsilyl-N-methyltrifluoroacetamide (c_final_: 6:7 (V/V); #394882, Sigma-Aldrich) with interspersed nitrogen-facilitated evaporation. Standards reflecting the natural ^13^C isotopomeric abundance pattern of the analytes of interest were similarly processed. Finally, the samples and standards were centrifuged (20,000 x *g*, 5 minutes, 4 °C) and the supernatant collected for analysis.

The prepared samples were separated and analysed in a gas chromatograph (Agilent Technologies; 7820A chromatograph; J&W GC column HP-5MS) coupled to a mass spectrometer (Agilent Technologies; 5977E). Metabolite abundances were integrated using the MassHunter software (Agilent Technologies). The metabolism-dependent isotopic enrichment of each metabolite of interest was estimated as previously described by Walls [66] and Biemann [67] by (a) correcting for the natural abundance of ^13^C-atoms in the ^13^C-labelling data by subtracting the mass spectrum of an unlabeled standard containing the metabolites of interest and (b) normalising the corrected abundance for a given isotopologue to the sum of corrected abundances of all isotopologues of the given metabolite. An unlabeled metabolite is registered as M in the mass spectrum, with a mass increase of 1 for each ^13^C-atom incorporated (M+1, M+2…, M+X). These are collectively called the isotopologues/isotopomers of a given metabolite.

Through glycolysis, [U-^13^C]D-glucose yield [U-^13^C]D-pyruvate, which through [1] oxidation by lactate dehydrogenase may yield [U-^13^C]L-lactate (lactate M+3), [2] amination by alanine transaminase may yield [U-^13^C]D-alanine (alanine M+3), or [3] oxidative decarboxylation by pyruvate dehydrogenase may yield [U-^13^C]acetyl-CoA. One forward metabolisation cycle of the latter yields M+2 isotopologue tricarboxylic acid (TCA) cycle intermediates and TCA-derived amino acid derivates. A 4-hour incubation time was selected based on preliminary experiments that demonstrated quantifiable isotopic enrichment of all glucose-derived TCA intermediates in the M+2 isotopologue pool, with other isotopologue pools of the TCA intermediates and related amino acids remaining relatively stable over time (S1 Fig). As such, the molecular ^13^C enrichment pattern of the M+3 isotopologue pool is reported for lactate and alanine, while the M+2 isotopologue pool ^13^C enrichment pattern is reported for all other metabolites.

### Targeted amino acid quantification by high-pressure liquid chromatography

Targeted amino acid quantification of the fibroblast was assessed by reverse-phase HPLC employing the 1260 Infinity (Agilent Technologies) as previously described [68].

Samples were prepared as described in the isotopic labelling experiments. MilliQ water-constituted cellular extracts were used for the analysis. Amino acid separation and detection were performed by precolumn o-phthalaldehyde online derivatisation and fluorescent detection (λ_ex_ = 338 nm, 10-nm bandwidth; λ_em_ = 390 nm, 20-nm bandwidth). A gradient elution with Mobile phase A (10 mM NaH_2_PO_4_, 10 mM Na_2_B4O_7_, 0.5 mM NaN_3_, pH 8.2) and mobile phase B (acetonitrile 45%: methanol 45%: H_2_O 10% V:V:V) was performed with a flow of 1.5 ml/min. Amino acid amounts were extrapolated from calibration curves based on external amino acid standards ranging from 5-1000 µM. Results were normalised to the protein amount of the cellular pellets determined by the BCA assay.

### Oxidative stress susceptibility assessment

Oxidative stress resilience of NTG and control fibroblasts was assessed using hydrogen peroxide (H_2_O_2_; #H1009, Sigma Aldrich) and a commercially available lactate dehydrogenase release assay kit (#11644793001, Merck).

Two days before each experiment, NTG and control fibroblasts were plated in 6-well plates at a density of 52,000 cells/cm^2^, and the culture medium was changed the following day. On the day of the experiment, fibroblasts were washed with 37 °C dPBS and incubated in culture medium supplemented with H_2_O_2_ (0 µM, 75 µM, 150 µM, and 300 µM) for 4, 24, and 48 hours. Experimental H_2_O_2_ doses were selected based on preliminary 24-hour dose-cytotoxicity screening experiments using the control K23 fibroblast cell line (S2 Fig). Negative media controls were processed simultaneously.

Incubation was terminated by collecting the media in prechilled Eppendorf tubes and then stored on ice until further processing. Fibroblasts were lysed immediately after media removal using ice-cold 1% PBST. Complete cell lysis was verified microscopically. Media and cell lysate were subsequently centrifuged at 20,000 × g for 10 minutes at 4°C, and the supernatants were collected and stored on ice. Media, cell lysate, and background supernatant were plated in triplicate in transparent 96-well plates on ice. Then, plates were transferred to regular tabletops and calibrated at RT for 10 min before assay initiation. Assay reactions were initiated by adding reaction mixture (1:45 V/V mixture of solution A and B, respectively) and terminated by adding 1 M hydrochloric acid after 12 min. Reactions were run shielded from light. Finally, the absorbance at wavelength λ = 490 nm on a spectrophotometer (Spectramax I3x, Molecular Devices; USA) was recorded. After background normalisation, relative extracellular lactate dehydrogenase release for a given sample was computed by normalising the media absorbance change at λ = 490 nm to the total absorbance change in the media and the cell lysate. Data for the H_2_O_2_-treated samples is presented relative to non-treated samples.

### Statistical analysis

Statistical data analysis was performed using the GraphPad Prism software version 10.4 (GraphPad Software, USA). The Shapiro-Wilk test was used to assess data distribution. Donor group characteristics were compared using a two-tailed unpaired t-test. Intergroup comparisons on continuous normally distributed data with multiple outcomes (e.g. data from seahorse, GCMS, HPLC and oxidative stress susceptibility experiments) were performed using two-tailed unpaired t-test with Welch’s correction and corrected for multiple comparisons using two-stage setup false discovery rate correction. Non-normally distributed data compared across multiple groups were analysed using the Kruskal-Wallis test with Dunn’s post hoc test for multiple comparisons. A significance level of 𝝰 = 0.05 and a false-negative rate of β = 0.2 were used for all statistical tests. p or q < 0.05 was defined as statistical significance. Specific data presentations are denoted for each analysis.

## Results

### Patient characteristics

Demographic and ophthalmological characteristics of the control and NTG donor groups are provided in Table 1. As expected, the NTG group exhibited a reduced perimetric mean deviation compared to the control group for both the right eye (MD OD) and the left eye (MD OS). Moreover, the control group demonstrated a slightly lower visual acuity (VA OD and OS) of both eyes compared to the NTG group, reflecting a differential cataract burden. No other group differences were shown.

**Table 1.** Patient baseline characteristics. **Table 1. Demographic and ophthalmic group characteristics of included patients.** Results are presented as medians with associated ranges. Data was analysed using two-tailed unpaired t-test. Statistical significance was defined as p < 0.05. BMI: Body Mass Index; IOP: Intraocular Pressure; MD: Mean Deviation; VA: Visual Acuity; SP: Spheric Correction; OD: Right Eye; OS: Left Eye; D: Dioptres.

| Variable | Control |  | NTG |  | P value |
| --- | --- | --- | --- | --- | --- |
|  | Median | Min-Max | Median | Min-Max |  |
| Age (years) | 69,5 | 67 - 78 | 69,5 | 67 - 75 | 0,82 |
| Height (cm) | 165 | 158 - 178 | 166 | 158 - 172 | 0,86 |
| **Weight (kg)** | 67,5 | 54 - 103 | 66,5 | 50 - 80 | 0,60 |
| **BMI (kg/m<sup>2</sup>)** | 25 | 21 – 33,51 | 24,1 | 20 – 27,04 | 0,52 |
| **IOP OD (mmHg)** | 13,5 | 13 - 16 | 14,5 | 12 - 18 | 0,62 |
| **IOP OS (mmHg)** | 13,5 | 13 - 17 | 14,5 | 12 - 18 | 0,76 |
| **MD OD (dB)** | **1,2** | **0,3 – 2,6** | **12,4** | **5,8 – 22,2** | **0,02** |
| **MD OS (dB)** | **0,45** | **-0,1 – 2,2** | **9,95** | **3,9 – 18,6** | **0,02** |
| **VA OD** | **0,57** | **0,32 - 1** | **1** | **0,9 – 1,25** | **0,04** |
| **VA OS** | 0,72 | 0,5 – 1,0 | 1 | 1 | 0,05 |
| **SP OD (D)** | 0,38 | -0,5 – 2,25 | 0,75 | -0,75 – 2,5 | 0,81 |
| **SP OS (D)** | 0,63 | 0,0 – 2,25 | 0,13 | -0,75 - 0,75 | 0,23 |

### Fibroblast culture purity is independent of donor origin

Fibroblast morphology and culture purity were assessed by phase-contrast microscopy and immunocytochemistry. All cultures exhibited typical morphology (e.g., elongated, spindle-shaped, or stellate mononuclear cells) and high purity, evaluated as the fraction of vimentin^+^/DAPI^+^ cells (Fig 2A-C).

**Fig 2.**
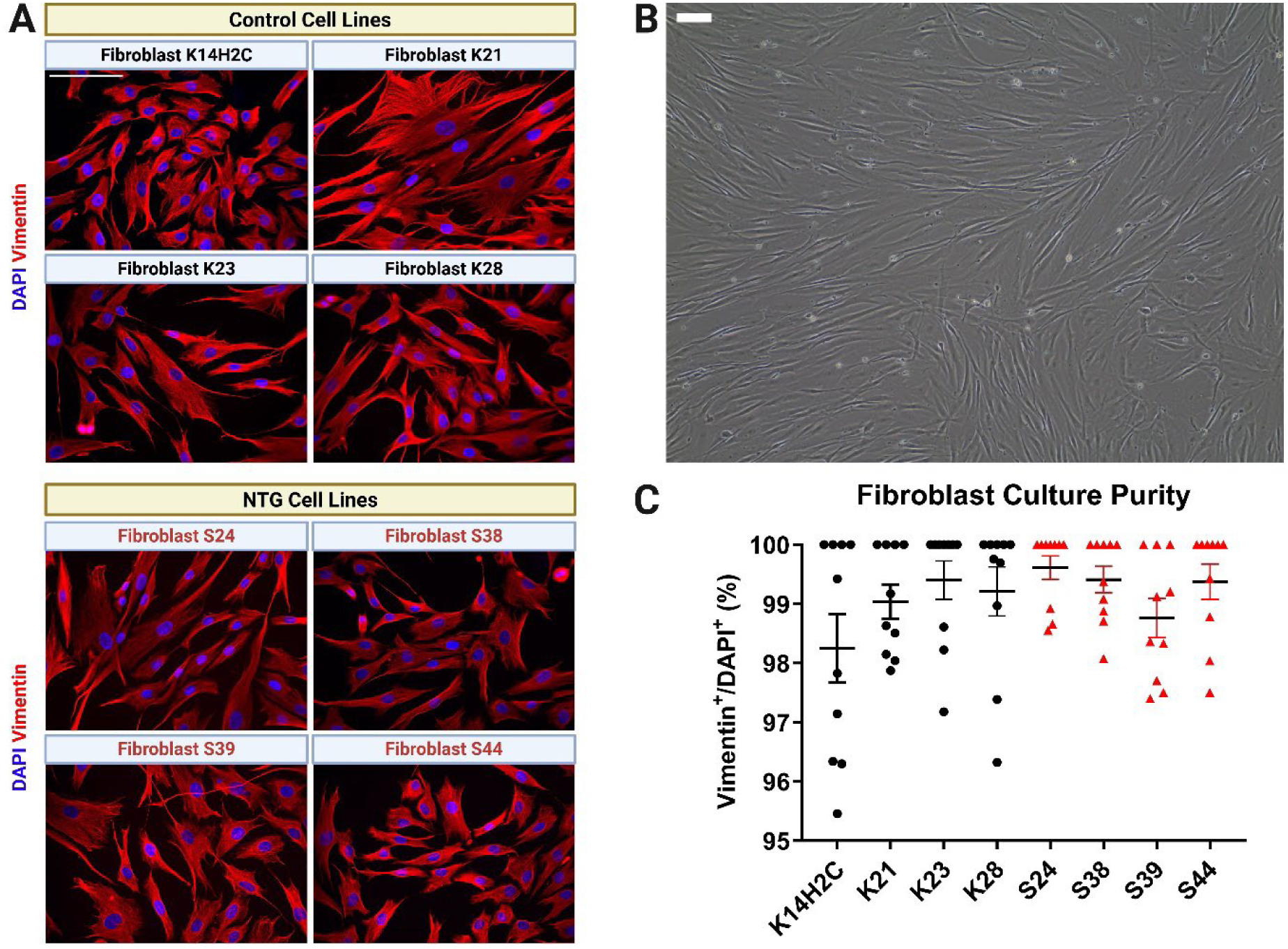
Fibroblast culture profiling. A: Immunocytochemical vimentin staining of the control (K14H2C, K21, K23, K28) and NTG (S24, S38, S39, S44) fibroblast cell lines. B: A representative phase contrast image of control fibroblast cell line K21 with similar morphological characteristics to other control and NTG fibroblast cell lines (not shown). C: Fibroblast culture purity comparisons using the cellular Vimentin^+^/DAPI^+^ culture fraction. Data is presented as means ± SEM for n = 10 technical replicates per cell line. Statistical comparisons were performed using the Kruskal-Wallis test with Dunn’s post hoc test. Statistical significance was defined as p < 0.05. Scale bars equal 100 µm. SEM: Standard error of the mean.

### Similar mitochondrial and glycolytic function of normal-tension glaucoma and control skin fibroblasts

Combined profiling of the mitochondrial and glycolytic function of skin fibroblasts from NTG and control patients was assessed using live-cell bioenergetics (Seahorse Analyser; Fig 3A-D). No statistically significant phenotypic differences could be demonstrated on any of the assessed functional parameters on mitochondrial respiration (Fig 3C) or anaerobic glycolytic function (Fig 3D).

**Fig 3.**
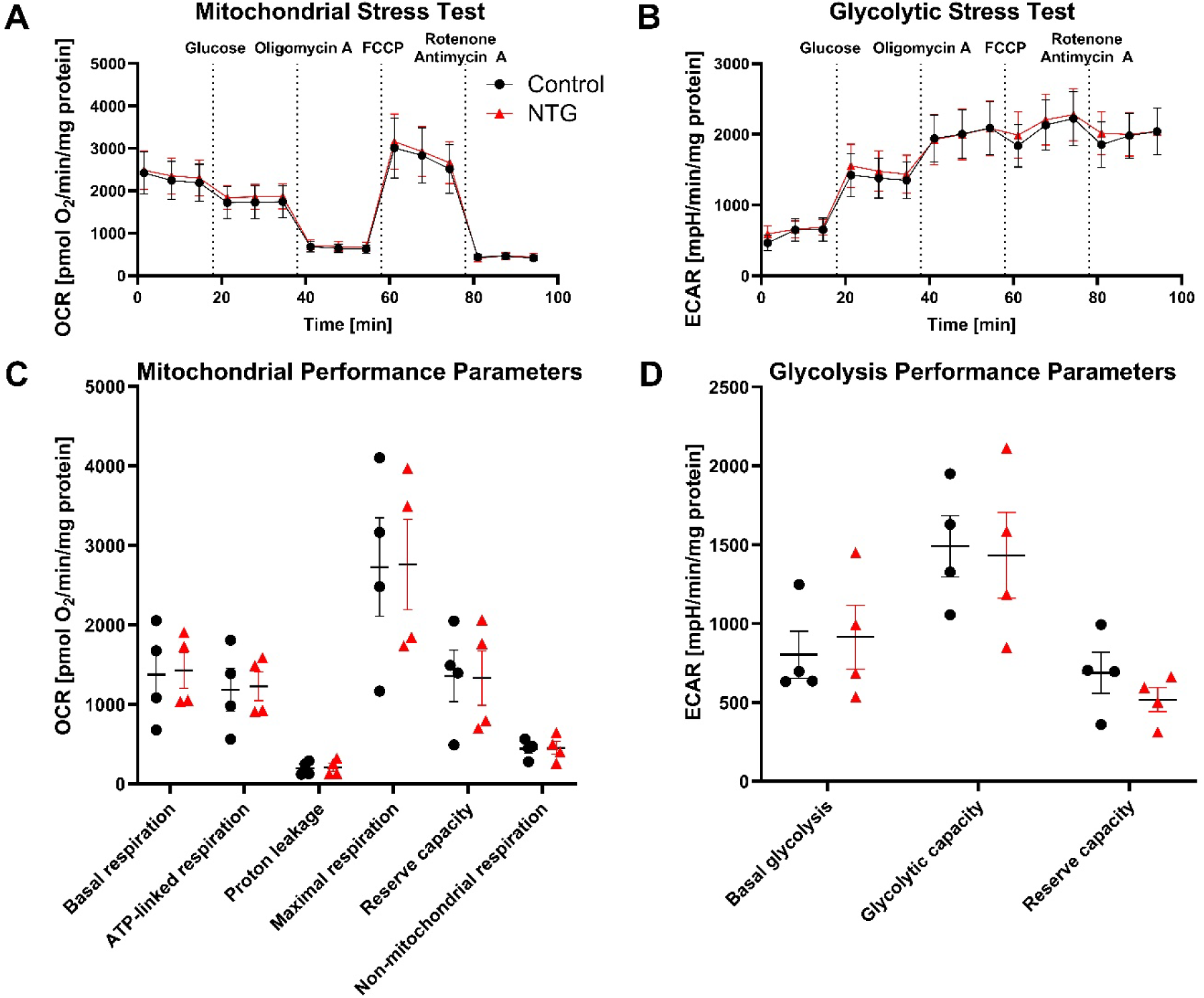
Bioenergetic profiles of normal-tension glaucoma (NTG) and control skin fibroblasts. (A) The mitochondrial and (B) glycolytic function of NTG and control fibroblasts with extrapolated (C) mitochondrial and (D) glycolytic function parameters are shown. No statistically significant phenotypic differences could be demonstrated on the assessed parameters. Results are represented as means ± SEM from three independent experiments, each with 10 technical replicates per cell line. Data was analysed using two-tailed unpaired t-test with Welch correction and corrected for multiple comparisons using two-stage setup FDR correction. q < 0,05 was defined as statistical significance. NTG: Normal-Tension Glaucoma; SEM: Standard Error of the Mean; OCR: Oxygen Consumption Rate; ECAR: Extracellular Acidification Rate; FDR: False discovery rate.

### Comparable glucose metabolism of NTG and control skin fibroblasts

Targeted metabolic profiling of NTG and control fibroblasts was performed using stable isotopic labelling with [U^13^C]glucose, GCMS, and HPLC. No statistically significant relative phenotypic differences in the ^13^C enrichment patterns of the M+3 isotopologues for lactate and alanine, nor the one-forward-cycle related M+2 TCA-cycle intermediates (citrate, α-ketoglutarate, succinate, malate) or associated amino acids (glutamate, glutamine, aspartate) could be demonstrated (Fig 4A). Moreover, no quantitative phenotypic differences could be shown in the amino acid profile (Fig 4B).

**Fig 4.**
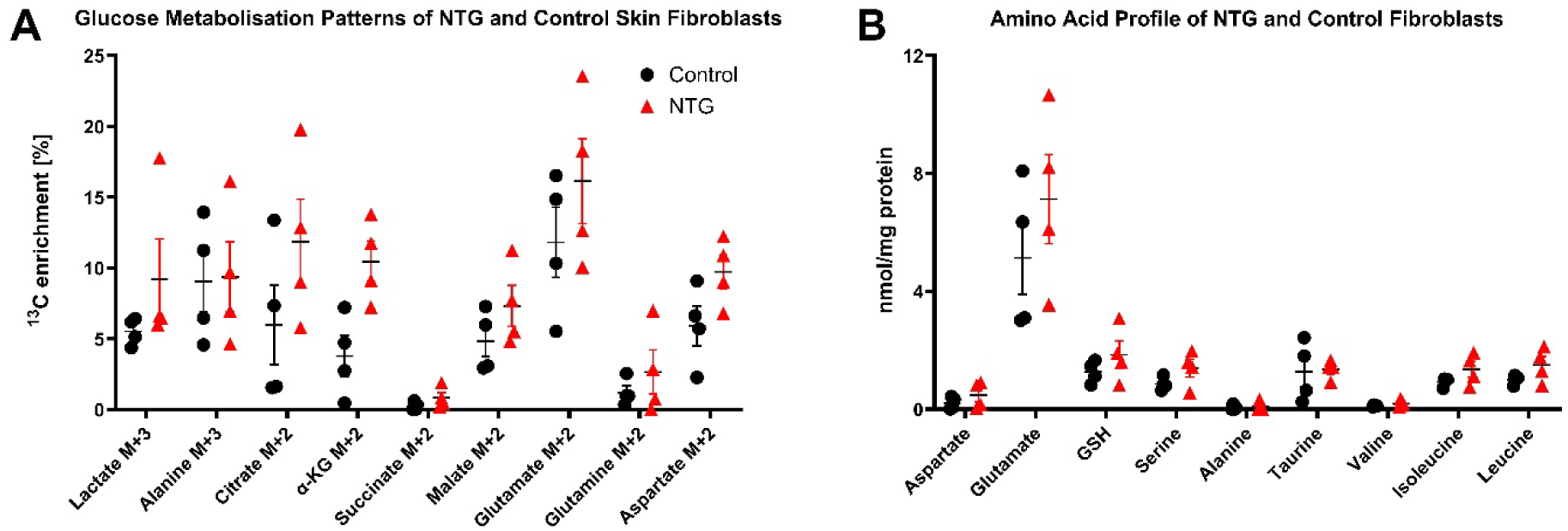
Relative glucose metabolism and absolute amino acid profiles of normal-tension glaucoma and control skin fibroblasts. A: Relative glucose metabolism patterns. B: Absolute amino acid content. No statistically significant phenotypic differences could be demonstrated on any of the assessed parameters. Results are represented as means ± SEM from three independent experiments with one technical replicate per cell line. Data was analysed using two-tailed unpaired t-test with Welch correction and corrected for multiple comparisons using two-stage setup FDR. q < 0,05 was defined as statistical significance. NTG: Normal-Tension Glaucoma; SEM: Standard Error of the Mean; FDR: False Discovery Rate.

### Similar oxidative stress susceptibility of NTG and control skin fibroblasts

The short- and long-term oxidative stress resistance of NTG and control skin fibroblasts was assessed using various concentrations of H_2_O_2_ (Fig 5). No statistically significant time-matched phenotypic differences in oxidative stress resiliency could be demonstrated for any of the assessed H_2_O_2_ concentrations.

**Fig 5.**
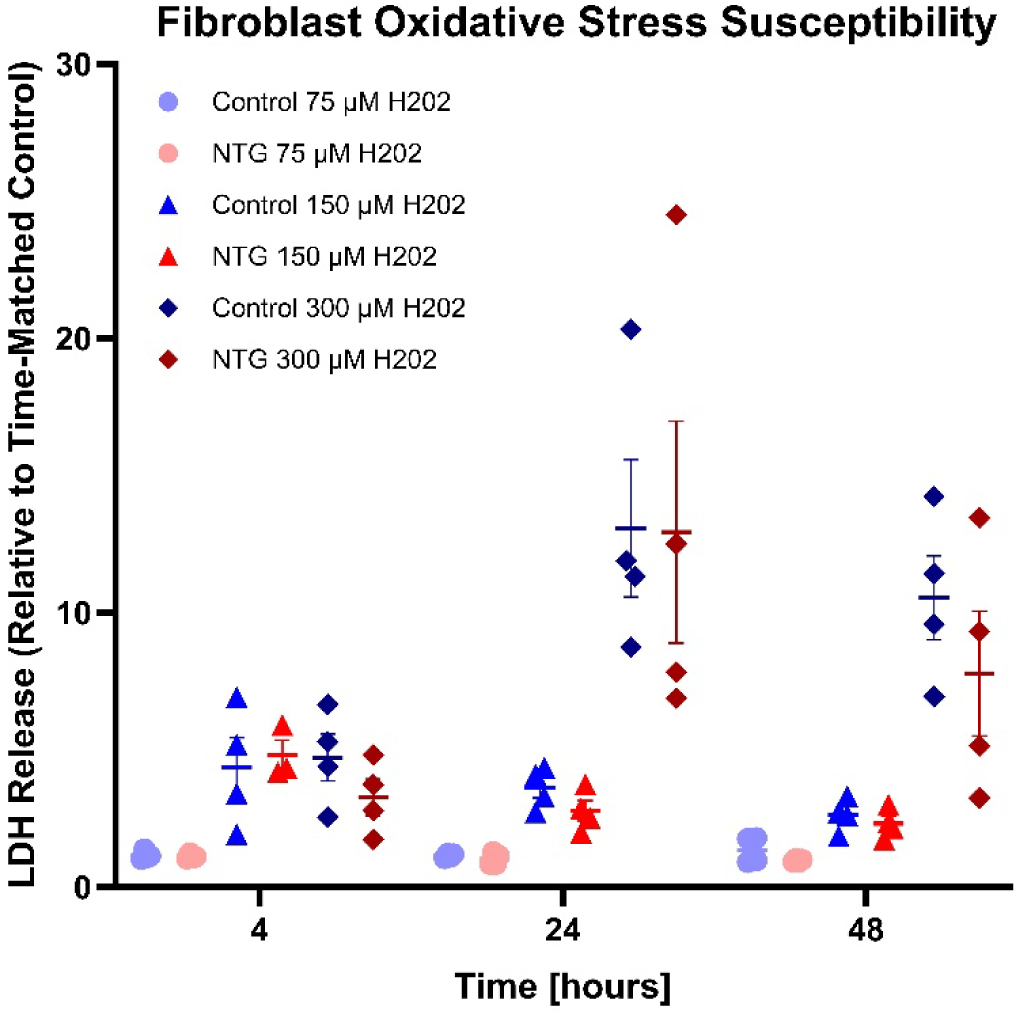
Concentration- and time-dependent oxidative stress resistance of NTG and control skin fibroblasts. Results are represented as means ± SEM from three independent experiments with one technical replicate per cell line. Data on time- and dose-matched phenotypic differences were compared using two-tailed unpaired t-test with Welch correction and corrected for multiple comparisons using two-stage setup FDR correction. q < 0,05 was defined as statistical significance. NTG: Normal-Tension Glaucoma; H2O2: Hydrogen Peroxide; SEM: Standard Error of the Mean.

## Discussion

In this study, we assess key metabolic and oxidative stress response signatures of NTG and control skin fibroblasts to elucidate cellular perturbations underlying vulnerability to IOP-independent glaucomatous neurodegeneration. Our study included assessments of mitochondrial and glycolytic function, focused glucose and amino acid metabolism profiling, and evaluation of exogenous oxidative stress resilience. No statistically significant differences were demonstrated between the donor groups in the assessed outcomes.

A capable bioenergetic metabolism, comprising anaerobic glycolysis and mitochondrial oxidative phosphorylation, is vital for maintaining retinal health and buffering against glaucomatous stress. [13]. The mitochondrion underlies several essential bioenergetic processes, and respiratory dysfunction has previously been demonstrated to be attenuated in several types of non-ocular cell types of POAG patients, especially NTG patients [29–32]. Additionally, pre-diagnostic systemic plasma deficiency of TCA-cycle substrates (citrate, 3-hydroxybutyrate, and acetate) and transferring molecules/cofactors (carnitine derivatives, biotin) increases the odds of a POAG [35]. Moreover, anaerobic glycolysis is a vital bioenergetic pathway for neurons and supporting cells of the central nervous system, including the retina [13, 69, 70]. In addition to ATP, lactate is an essential byproduct due to its neuroprotective properties mediated through glia-mediated metabolic support [71–74] and survival signalling [75]. Analogously, systemic evidence of diminished anaerobic glycolysis has likewise been obtained from POAG patients [34, 35, 74]. As such, some POAG patients exhibit reduced plasma levels of pyruvate [35] and lactate [34, 35, 74]. Overall, any sort of disturbance to substrate availability, glycolysis, or oxidative phosphorylation is likely to enact detrimental effects on RGC health, as systemic and glia-mediated metabolic buffering seems to be the primary mechanisms to protect RGCs during stress [22, 35, 71–73, 76]. Concerning our findings showing similar mitochondrial respiratory and glycolytic function of NTG fibroblasts evaluated by the Seahorse assay and by targeted investigation of glucose metabolisation using stable isotopic labelling and GCMS-metabolomics, our study does not support the presence of intrinsic bioenergetic defects in these processes in skin fibroblasts from NTG patients relative to controls. Future studies assessing additional aspects of mitochondrial function and dynamics [77], processes that likewise exhibit disturbances on a genetic background in other optic neuropathies [42, 78], are warranted.

Metabolomic analysis of several fluid compartments in POAG patients demonstrates differential changes in several key amino acid, carbohydrate, and lipid pathways [22, 35], including the aqueous humour, tears, plasma/serum, and optic nerve. Affected areas include, among others, the metabolism of glycine, serine, threonine, glutathione, taurine, alanine, aspartate, and glutamate [22, 35]. Metabolic disturbance in these systems could have detrimental consequences due to the widespread implications of these processes. Several of these shuttle/support systems (the aspartate-malate shuttle, the glutamine-glutamate cycle, the lactate shuttle) are responsible for supporting neuronal metabolism by maintaining cytosolic NAD, fuelling the ETC with cytosolic-generated electrons, and providing substrates for the TCA-cycle [13, 70]. Glutathione is vital for maintaining cellular NAD^+^/NADP^+^ levels, thereby buffering cellular antioxidant capacity and regulating metabolic flow, which is especially important in the retina [79]. Glycine, serine, and threonine are vital for glutathione biosynthesis, DNA repair, methylation, the synthesis of several essential cofactors (e.g., folate), and cell survival signalling [80]. Reduced levels have been shown to promote retinal degeneration in animal models of retinopathy [80]. Taurine is the most abundant retinal amino acid and possesses several retinoprotective properties, including anti-oxidation, anti-apoptotic signalling, osmoregulation, and counteracting glutamate-induced excitotoxicity [81]. With respect to our findings showing similar relative glucose-dependent biogenesis patterns of a lactate, alanine, citrate, α-ketoglutarate, succinate, malate, glutamate, glutamate, glutamine, and aspartate as well as no absolute quantity differences in aspartate, glutamate, glutathione, serine, alanine, taurine, valine, isoleucine, and leucine cellular content, this current study does not support the presence of intrinsic bioenergetic disturbance on the assessed parameters in NTG fibroblasts compared to controls. Future studies performing an extended analysis of the metabolome are warranted.

Oxidative stress is a hallmark of glaucoma and may be a consequence of the underlying pathogenesis of glaucoma [50, 51]. Hence, mitochondrial dysfunction, vascular dysregulation, IOP fluctuations, excitotoxicity, and reactive immunity all converge on oxidative stress, but can likewise be a downstream consequence thereof [82–84]. Thus, an intrinsically compromised cellular/non-cellular antioxidation system or increased endogenous production of oxidants will likely influence glaucoma susceptibility. In this regard, POAG patients exhibit diminished plasma antioxidant capacity [54–57], while also displaying increased features of oxidative stress damage [56, 57]. To investigate the presence of differential cellular resistance to oxidative stress, NTG and control fibroblasts were exposed to H_2_O_2_ for different periods, and the cytotoxicity was assessed. No intergroup difference could be demonstrated, indicating no differences in resistance to H_2_O_2_-induced oxidative stress [85]. In this context, complementary future studies focusing on response differences from extended periods of (intermittent) low-grade oxidative stress, such as hyperoxia exposure, which may more accurately mimic in vivo NTG conditions (i.e., microvascular dropout) [86], are warranted. Moreover, no group differences in cellular non-mitochondrial oxygen consumption rates were observed, suggesting a similar baseline production of non-mitochondrial reactive oxygen species. However, direct measures of mitochondrial and non-mitochondrial ROS should be performed for definitive verification.

The findings of the current study should be interpreted in consideration of its limitations. Despite including pure clinical NTG phenotypes, the present study is likely statistically underpowered, considering the demonstration of minor phenotypic effect differences, which may be highly relevant in glaucoma [32, 47, 49, 87]. Moreover, as several endotypes converge on the NTG phenotype, proper genetic and transcriptomic profiling should be used to guide and stratify functional profiling outcomes per patient case [47–49]. In this context, tissue- and cell-specific differences resulting from differential gene expression should be considered and warrant investigation in future studies [13, 58]. This is needed to identify valid endotype-specific bio-and predictive markers for POAG susceptibility, risk profiling, and visual prognosis [32]. Despite obtaining pure fibroblast cultures for all donors while attempting to limit bias arising from differences in anatomical origin and donor differences, fibroblast heterogeneity resulting from exogenous stimuli and lifestyle-related differences is another confounding factor that enhances interindividual variability [88, 89]. Finally, as the present study focuses solely on females, future studies examining systemic predispositions in male NTG donors are warranted.

## Acknowledgements

We want to acknowledge Charlotte Taul, laboratory manager at the Department of Drug Design and Pharmacology at the University of Copenhagen (Denmark), for her assistance with maintaining fibroblast cell cultures.

## Author contributions

AVS, RV, BIA, KF, and MK contextualised the study. AVS, SS, and MK acquired funding. AS, SS, and MK recruited patients. AVS and KSD performed the experiments. AVS and KSD performed data analysis. All authors interpreted the findings. AVS drafted the manuscript. All authors reviewed and approved the final version submitted for publication.

## Data availability

Due to legal confidentiality, primary patient data is not available. Experimental data is available at https://osf.io/cbhn8/?view_only=6364e0ab6e064d1a9d6dac988a100c34.

