## Supplemental Figures for "Inferring systemic metabolic and oxidative stress susceptibility in normal-tension glaucoma through targeted skin fibroblast analysis"

#### Figure S1

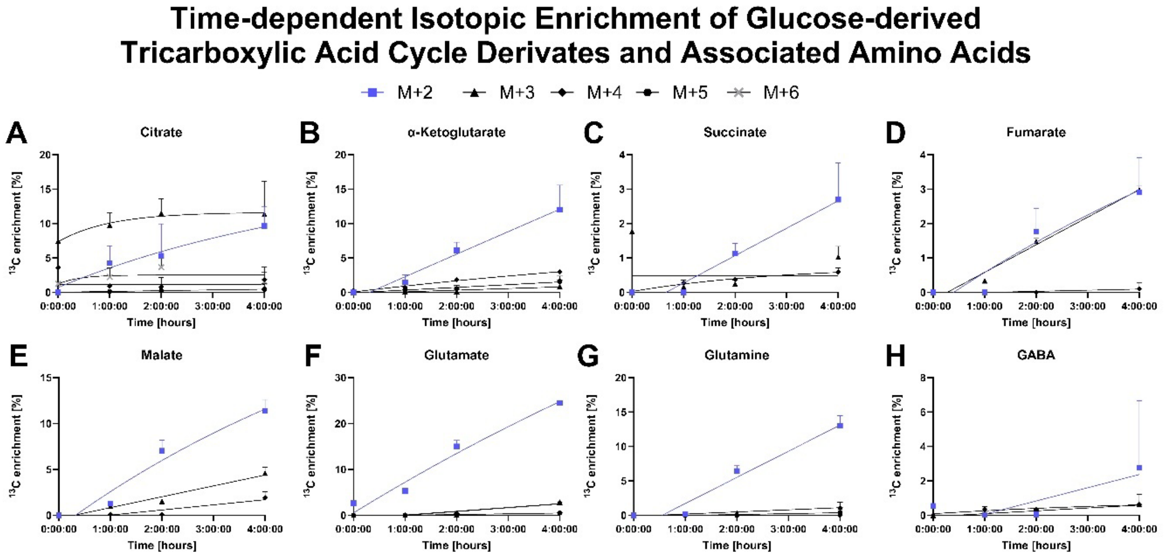

*__Figure S1__. $^{13}$C-enrichment [%] of the M+1 to M+6 isotopologues of selected TCA cycle intermediates (A-E) and derived amino acids (F-H) following incubation with 6 mM [U-$^{13}$C]glucose for 0-4 hours using the control K23 skin fibroblast cell line. The preliminary test on one control fibroblast cell line was justified on the preliminary thesis of distinctive donor group differences with the control cell lines exhibiting enhanced oxidative stress resistance compared to NTG cell lines. Results are presented as mean ± SEM for three technical replicates for one preliminary experiment. Non-linear, one-phase association regression curves have been fitted to each dataset to illustrate the time-dependent increase in the $^{13}$C-labelling pattern of each isotopologue for the assessed metabolites. Intraisotopomeric, time-dependent differences in the $^{13}$C-labelling pattern were evaluated by Ordinary one-way ANOVA and Tukey's multiple comparison post hoc test with a two-sided significance of* $\alpha$ *= 0.05. Statistical data are not displayed on the plot. SEM: Standard Error of the Mean.*

### Figure S2

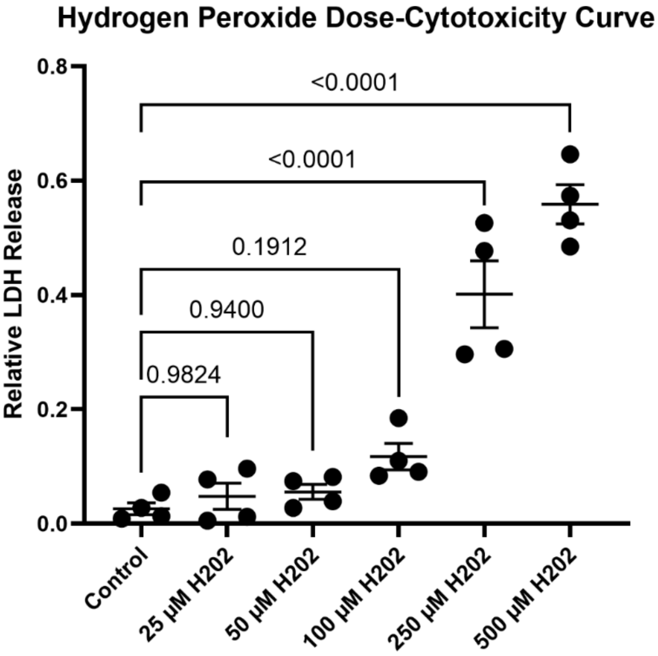

***Figure S2:*** 24-hour $H_2O_2$ dose-cytotoxicity curve for the control K23 fibroblast cell line. The preliminary test on one control fibroblast cell line was justified on the preliminary thesis of distinctive donor group differences with the control cell lines exhibiting enhanced oxidative stress resistance compared to NTG cell lines. The fibroblast cell line was incubated for 24 hours with different concentrations of $H_2O_2$ *(0, 25, 50, 100, 250 and 500 μM) and the relative LDH release quantified using the LDH release assay. Results are presented as mean ± SEM from one preliminary experiment using four technical replicates per dosage. Data was analysed using Ordinary one-way ANOVA, followed by Dunnett's multiple comparisons post hoc test. $p < 0.05$ defined statistical significance.* $H_2O_2$: Hydrogen Peroxide; *LDH: Lactate Dehydrogenase.*
